# Endogenous amdoparvovirus-related elements reveal insights into the biology and evolution of vertebrate parvoviruses

**DOI:** 10.1101/224584

**Authors:** Judit J Pénzes, Soledad Marsile-Medun, Mavis Agbandje-McKenna, Robert James Gifford

## Abstract

Amdoparvoviruses (family *Parvoviridae:* genus *Amdoparvovirus*) infect carnivores, and are a major cause of morbidity and mortality in farmed animals. In this study, we systematically screened animal genomes to identify PVe disclosing a high degree of similarity to amdoparvoviruses, and investigated their genomic, phylogenetic and protein structural features. We report the first examples of full-length, amdoparvovirus-derived PVe in the genome of the Transcaucasian mole vole (*Ellobius lutescens*). Furthermore, we identify four further PVe in mammal and reptile genomes that are intermediate between amdoparvoviruses and their sister genus (*Protoparvovirus*) in terms of their phylogenetic placement and genomic features. In particular, we identify a genome-length PVe in the genome of a pit viper (*Protobothrops mucrosquamatus*) that is more like a protoparvovirus than an amdoparvovirus in terms of its phylogenetic placement and the structural features of its capsid protein (as revealed by homology modeling), yet exhibits characteristically amdoparvovirus-like genome features including: (i) a putative middle ORF gene; (ii) a capsid gene that lacks a phospholipase A2 (PLA2) domain; (iii) a genome structure consistent with an amdoparvovirus-like mechanism of capsid gene expression. Our findings indicate that amdoparvovirus host range has extended to rodents in the past, and that parvovirus lineages possessing a mixture of proto- and amdoparvovirus-like characteristics have circulated in the past. In addition, we show that PVe in the mole vole and pit viper encode intact, expressible replicase genes that have potentially been co-opted or exapted in these host species.

## Introduction

Parvoviruses (family *Parvoviridae*) are small, single-stranded DNA viruses that infect vertebrate (subfamily *Parvovirinae*) and invertebrate (subfamily *Densovirinae*) hosts. The small (4-6 kb) genome is encompassed by characteristic terminal palindromic repeats, which form hairpin-like secondary structures characteristic for each genus (Cotmore et al. 2014; Tijssen et al. 2011). Despite exhibiting a low level of sequence similarity, parvovirus genomes are highly conserved in overall structure, containing two large gene cassettes responsible for encoding the non-structural (NS) and the structural (VP) proteins. The N-terminal region of the minor capsid protein VP1 includes a highly conserved phospholypase A2 (PLA2) motif that is required for escape from the endosomal compartments after entering the host cell (Zadori et al. 2001). The parvovirus capsid is icosahedral with a T=1 symmetry, displaying a jelly roll fold of conserved β-sheets linked by variable surface loops, designated variable region (VR) I to IX (M.S. Chapman and Agbandje-McKenna 2006).

Endogenous parvoviral elements (PVe) are parvovirus sequences that are thought to have been generated when DNA sequences derived from viruses were incorporated into the germline of the ancestral host species (via infection of germline cells), such that they were subsequently inherited as host alleles (Holmes 2011). These sequences provide a useful source of information about the biology and evolution of ancient parvoviruses (Feschotte and Gilbert 2012; Holmes 2011). For example, the identification of orthologous PVe in the genomes of distantly related mammalian species demonstrates an association between parvoviruses and mammals that extends over tens of millions of years (Belyi et al. 2010; Kapoor et al. 2010; Katzourakis and Gifford 2010; Smith et al. 2016).

*Amdoparvovirus* is a recently defined genus in the family *Parvoviridae* (Cotmore et al. 2014). The type species was originally called Aleutian mink disease virus (AMDV) – hence the genus name. However, AMDV is now considered to represent a variant of the renamed species *Carnivore amdoparvovirus 1* (Cotmore et al. 2014). AMDV causes an immune complex-associated progressive syndrome in American mink (*Neovison vison*) called Aleutian disease (AD) or plasmacytosis (Bloom et al. 1994) which is considered to be one of the most important infectious diseases affecting farm-raised mink (Canuti et al. 2015; Canuti et al. 2016). AMDV infection is known to be widespread in wild mink as well as in farmed animals (Canuti et al. 2015), and related amdoparvoviruses have been identified in other carnivore species, including raccoon dogs, foxes, skunks, and red pandas (Alex et al. 2018; LaDouceur et al. 2015; Li et al. 2011; Pennick et al. 2007; Shao et al. 2014). Findings from metagenomic studies suggest that amdoparvoviruses may infect a broader range of mammalian orders (Bexfield and Kellam 2011), but this has yet to be fully demonstrated.

Phylogenetic studies support a common evolutionary origin for amdoparvoviruses and protoparvoviruses (genus *Protoparvovirus*) (Cotmore et al. 2014). Protoparvoviruses infect a wide range of mammalian hosts, encompassing several distinct mammalian orders (Sasaki et al. 2015; Tijssen et al. 2011). For example, rodent protoparvoviruses are known for their oncolytic properties (Marchini et al. 2015), while carnivore and ungulate protoparvoviruses are significant pathogens of domestic pets and livestock (Hueffer and Parrish 2003; Kailasan et al. 2015; Meszaros et al. 2017). Although they are relatively closely related, amdoparvoviruses and protoparvoviruses are distinguished by certain features of their genomes and replication strategies. In particular, amdoparvovirus mRNAs are transcribed from one single upstream promoter and are polyadenylated at two distinct polyadenylation signals. To provide the VP1 encoding transcript, an intron is spliced out, leaving a short, three amino acid-encoding exon leader sequence (transcribed from a short, upstream ORF) positioned in-frame with the VP ORF (Qiu et al. 2006). In contrast, the NS and VP-encoding mRNAs of protoparvoviruses are transcribed from two different promoters, being polyadenylated at a mutual polyadenylation signal close to the 3’ end of the genome. Although splicing has been reported in the protoparvovirus VP-encoding mRNA, this always occurs within the VP ORF itself (Tijssen et al. 2011). Amdoparvoviruses are also unique within the *Parvovirinae* in lacking a PLA2 domain in their VP unique region (VP1u) (Cotmore et al. 2014; Zadori et al. 2001).

In this study, we performed a systematic screen of 688 animal genomes to identify PVe disclosing similarity to amdoparvoviruses. We identify and characterise six such PVe, examining their genomic, phylogenetic and protein structural characteristics.

## Methods

### in silico genome screening

We used the database-integrated genome screening (DIGS) tool (Zhu et al. 2018) to screen while genome sequence (WGS) assemblies for PVe. The DIGS tool provides a framework for similarity-search based genome screening. It uses the basic local alignment search tool (BLAST) program (Altschul et al. 1997) to systematically screen WGS files for sequences matching to a nucleotide or peptide ‘probe’. Sequences that disclose above-threshold similarity to the probe are extracted and classified, with results being captured in a MySQL relational database (Axmark and Widenius 2015). To identify PVe, we used parvovirus peptide sequences to screen all 362 vertebrate genome assemblies available in the NCBI whole genome sequence (WGS) database as of the 15^th^ December 2017. Sequences that produced statistically significant matches to these probes were extracted and classified by BLAST-based comparison to a set of reference peptide sequences selected to represent the broad range of diversity in subfamily *Parvovirinae.*

### Sequence analyses

The characterization and annotation of PVe was performed using Artemis Genome Browser (Carver et al. 2012). The putative peptide sequences of PVe were inferred and aligned with NS and VP sequences of representative amdoparvoviruses and protoparvoviruses using MUSCLE (Edgar 2004), PAL2NAL (Suyama et al. 2006) and T-coffee Expresso (Armougom et al. 2006). Phylogenies were reconstructed from amino acid (aa) alignments incorporating structural data (at least one high-resolution structure from all available genera) using maximum likelihood as implemented in PhyML-3.1, and 1000 bootstrap replicates (Guindon et al. 2010). Model selection was carried out by ProTest (Abascal et al. 2005) selecting the RtEV (NS) and the LG (VP) protein substitution models.

To detect structural homology, we applied the pGenTHREADER and pDomTHREADER algorithms of the PSIPRED Protein Sequence Analysis Workbench (Lobley et al. 2009). The selected PDB structures were applied as templates for homology modeling, carried out by SWISS-MODEL (Biasini et al. 2014). Polymers of the acquired capsid monomer models were constructed by the Oligomer Generator feature of the Viper web database (http://viperdb.scripps.edu/) (Carrillo-Tripp et al. 2009). The generated polymers were rendered as well as ribbon diagrams compared using PYMOL (Schrödinger)

## RESULTS

### Identification and characterisation of PVe

We screened whole genome sequence (WGS) assemblies of 688 animal species for PVe disclosing a high degree of homology to amdoparvoviruses. Similarity searches using the replicase (NS) and capsid (VP) proteins of AMDV identified six such PVe (**Table 1**). These sequences were identified in five distinct vertebrate species, including a reptile – the spotted pit viper - in addition to four mammals. The mammals included three placental species (a rodent and two afrotherians) and one marsupial.

**Table 1.**
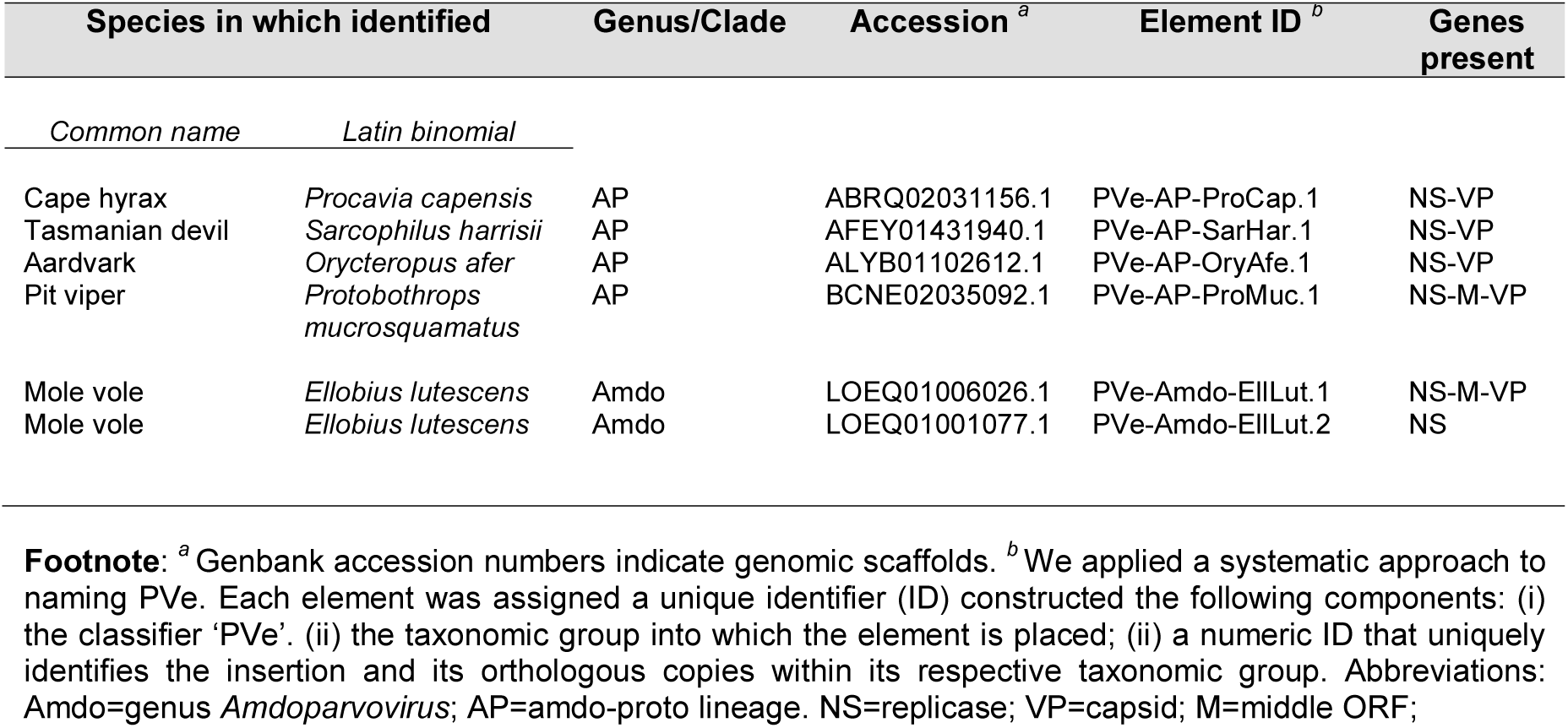
PVe characterised in this study.

| Species in which identified |  | Genus/Clade | Accession <sup>a</sup> | Element ID <sup>b</sup> | Genes present |
| --- | --- | --- | --- | --- | --- |
| *Common name* | *Latin binomial* |  |  |  |  |
| Cape hyrax | *Procavia capensis* | AP | ABRQ02031156.1 | PVe-AP-ProCap.1 | NS-VP |
| Tasmanian devil | *Sarcophilus harrisii* | AP | AFEY01431940.1 | PVe-AP-SarHar.1 | NS-VP |
| Aardvark | *Orycteropus afer* | AP | ALYB01102612.1 | PVe-AP-OryAfe.1 | NS-VP |
| Pit viper | *Protobothrops mucrosquamatus* | AP | BCNE02035092.1 | PVe-AP-ProMuc.1 | NS-M-VP |
| Mole vole | *Ellobius lutescens* | Amdo | LOEQ01006026.1 | PVe-Amdo-EllLut.1 | NS-M-VP |
| Mole vole | *Ellobius lutescens* | Amdo | LOEQ01001077.1 | PVe-Amdo-EllLut.2 | NS |

**Footnote:** <sup>a</sup> Genbank accession numbers indicate genomic scaffolds. <sup>b</sup> We applied a systematic approach to naming PVe. Each element was assigned a unique identifier (ID) constructed the following components: (i) the classifier 'PVe'. (ii) the taxonomic group into which the element is placed; (ii) a numeric ID that uniquely identifies the insertion and its orthologous copies within its respective taxonomic group. Abbreviations: Amdo=genus *Amdoparvovirus*; AP=amdo-proto lineage. NS=replicase; VP=capsid; M=middle ORF;

In all cases, sequences were identified in contigs that were orders of magnitude larger than a parvovirus genome, and it was clear they represented PVe as opposed to contaminating virus (see below). All six loci were examined using sequence comparison tools to determine their genomic structure relative to reference viruses, and identify the locations of other genomic features, such as promoters, polyadenylation signals, and transposable element insertions (**Figure 1**). Comparisons of genomic sequences flanking PVe established that all six are present at distinct locations, and were generated in distinct germline incorporation events. To infer the evolutionary relationships of these elements to contemporary parvoviruses we reconstructed maximum likelihood (ML) phylogenies using conserved regions of their putative NS and VP peptide sequences (**Figure 2**). The genomic and phylogenetic characteristics of all six elements are described below.

**Figure 1.**
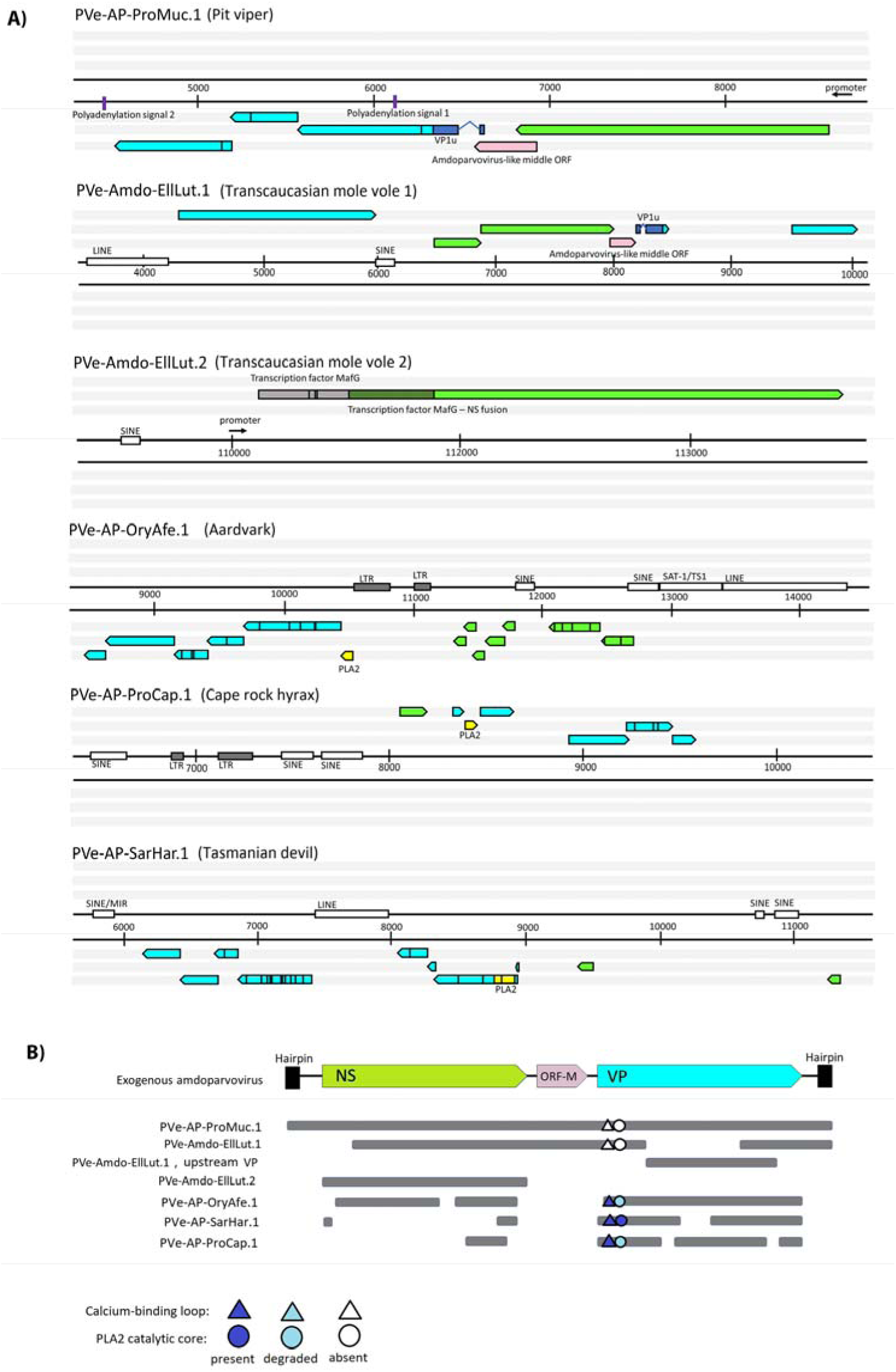
Genomic organization of six amdoparvovirus-like PVe. **Panel (a)**: Genomic structure of PVe loci, showing features identified in all six frames. Regions of homology to parvovirus proteins are indicated as arrows (NS in green, VP in cyan). Stop codons are indicated by vertical black lines, putative promoters with small black arrows. The characteristic middle (M) ORF homologs of amdoparvoviruses are shown in pink. In the EilLut.2 element, dark green represents the potentially expressed, NS-fused region of the MafG transcription factor. The remaining portion of the MafG pseudogene is shown in grey. **Panel (b)**: Genomic organisation of PVe, shown in relation to a representative amdoparvovirus genome. Abbreviations: NS – non-structural protein; VP – capsid protein; VP1u – VP1 unique region; LINE – long interspaced nuclear element; SINE – short interspaced nuclear element; LTR – long terminal repeat; PLA2 – phospholipase A2 domain.

**Figure 2.**
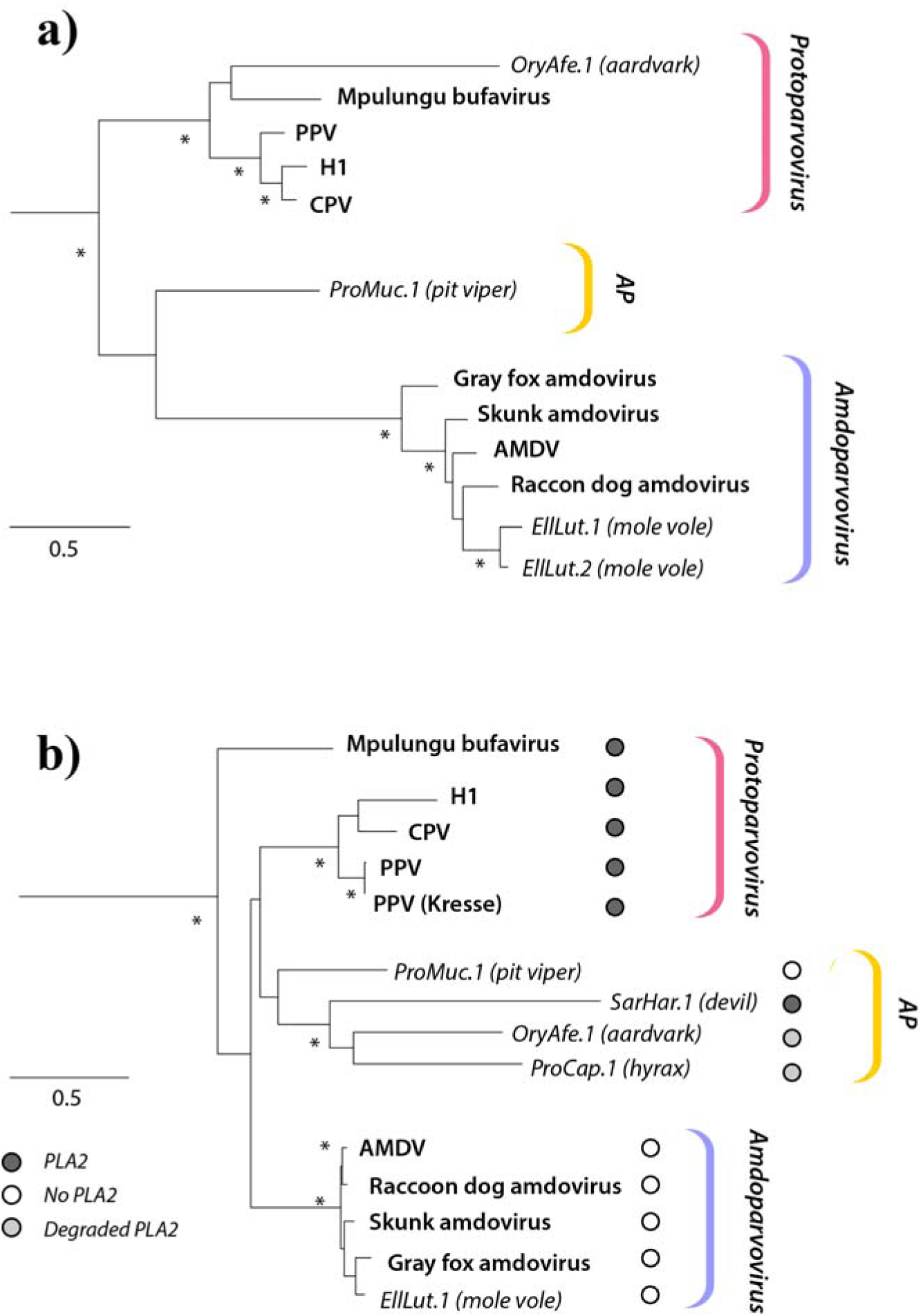
Maximum likelihood (ML) phylogenies of amdoparvoviruses, protoparvovirus and PVe. Phylogenies based on NS (**panel a**) and VP (**panel b**) peptide sequences. Viral taxa are shown in bold text. The taxa names of endogenous parvoviral elements (PVe) are shown in italics. Brackets to the right indicate viral genera (*Amdoparvovirus, Protoparvovirus*) and PVe clades. Asterisks indicate nodes with bootstrap support >90%. The scale bar shows genetic distance in substitutions per site. Abbreviations: AMDV=Aleutian mink disease virus; CPV=Canine parvovirus; PPV=Porcine parvovirus; MVM=Minute virus of mice; H1=H-1 parvovirus; AP=Proto-Amdo clade of PVe. Details of PVe examined here are contained in **Table 1. Table S1** contains the accession numbers and other details of amdoparvovirus and protoparvovirus reference sequences.

### Amdoparvovirus-derived PVe in a rodent genome

We identified two PVe derived from amdoparvoviruses in the genome of the Transcaucasian mole vole (*Ellobius lutescens*) (**Table 1**). ML phylogenes reconstructed using the putative peptide sequences encoded by these elements were used to reconstruct showed that both mole vole elements were closely related to one another, and grouped robustly within the clade defined by exogenous amdoparvoviruses (**Figure 2**). The first (PVe-Amdo-EllLut.1) spanned a near complete genome containing both the NS and VP genes, while the second (PVe-Amdo-EllLut.2) spanned the majority of the NS gene, with no identifiable VP present (see **Figure 1**). These elements are hereafter referred to as EllLut.1 and EllLut.2, respectively.

EllLut.1 is integrated into a locus that is homologous to mouse chromosome 12. This element is derived from genome-length nucleic acid, and contains a putative 3’ untranslated region (UTR) that exhibits homology to the the 3’UTR of AMDV and contains inverted repeats capable of folding into a stem loop structure (**Figure S1**).

The putative NS ORF of EllLut.1 has gaps relative to AMDV: the N-terminal region of NS is absent up to residue 28G. The NS ORF is flanked on either side by regions of VP homology (see **Figure 1**). A partial VP ORF could be identified downstream of the NS gene, corresponding to the VP1u and the VP2 N-terminal, as well as nucleotides encoding the last 173 amino acids (aa) of the C-terminus. However, a large part of the ORF is missing due to an assembly gap. A region of VP homology - spanning residues 59 to 600 - could be identified upstream of the NS ORF, encompassed by LINE and SINE elements. Only 60 aa of its derived protein sequence overlap with the downstream, partial VP ORF, indicating that the intervening assembly gap region may correspond to the missing portion of the VP gene.

Splicing of a putative intron sequence may position three residues of the short, 23-aa-long upstream adjacent ORF in-frame with the VP ORF, consistent with the typical VP1u transcription pattern of amdoparvoviruses (Qiu et al. 2006). The putative VP ORF encoded by EllLut.1 has a gap relative to the AMDV VP that spans most of the 5’ region of the gene. Frameshifting mutations are present in the NS genes of both elements, and the VP pseudogenes of EllLut.1 (**Figure 1**).

Amdoparvovirus genomes encode a short middle ORF (M-ORF) of unknown function between the two major ORFs (NS and VP) (Bloom et al. 1988; Gottschalck et al. 1994; Li et al. 2011). As shown in **Figure 1**, a region of potentially protein-coding sequence that corresponds to the M-ORF of AMDV is present in the EllLut.1 element. A methionine (M) residue that might represent the start codon of an M-ORF gene product could not be identified. However, this is also the case for several exogenous amdoparvovirus isolates (Gottschalck et al. 1994; Li et al. 2011).

The EllLut.2 element comprises the NS gene alone (**Figures 1**). This element is integrated into a locus immediately adjacent to the sequences encoding the MAF BZIP transcription factor G (MAFG) gene, which in the mouse genome is located in the 11qE2 region of chromosome 11. The structure of the EllLut.2 element indicates it was derived from an NS-encoding mRNA that was reverse transcribed and integrated into the nuclear genome of an ancestral germline cell. The otherwise intact NS gene lacks a methionine start codon and 5’ UTR. However, immediately upstream of the three stop codons disrupting the MAFG gene, a conventional ATG start codon was identified that could provide translation initiation to express a MAFG-NS fusion product (**Figure 1**). The identification of a potential promoter sequence downstream of the MAFG gene supports the existence of such a fusion protein, as does the strong Kozak translational context of the above mentioned start codon.

We identified empty integration sites in the *E. talpinus* genome at the loci where the EllLut.1 and EllLut.2 elements are integrated in *E. lutescens.* This indicates that both elements were integrated into the *E. lutescens* germline after these two species diverged ~10 million years ago (MYA) (Fabre et al. 2012; Pisano et al. 2015). The genomes of two *E.lutescens* individuals have been generated (genomic DNA was obtained from the livers of both a male and a female individual). Both PVe were present in both individuals. However, the EllLut.2 element in the female animal had a 13-14 bp deletion relative to the one in the male.

### A PVe in the pit viper genome with amdoparvoviral and protoparvoviral characteristics

We identified a PVe sequence in the genome of the spotted pit viper (*Protobothrops mucrosquamatus*), which we labelled PVe-AP-ProMuc.1 (ProMuc.1). This element, which encoded a nearly complete parvovirus genome, was ~4.5 kb in length and was integrated in reverse orientation (i.e. preserving the presumed original negative orientation of the virus genome). The putative genome structure comprised two major ORFs and a minor ORF as well as a clearly-identifiable and potentially functional downstream promoter (**Figure 1**). Furthermore, two polyadenylation signals could be identified, as well as partial palindromic repeats resembling the amdoparvoviral hairpin structures in the expected positions upstream and downstream of the two ORFs (**Figure 1, Figure S1**).

The first major ORF exhibited a relatively high degree of aa identity to the AMDV NS protein (35% with no deletions). The putative peptide gene product clustered as an outgroup to a clade containing the mole vole PVe and exogenous amdoparvoviruses (**Figure 2a**), but only with weak bootstrap support. The second major ORF, which was disrupted by several nonsense mutations (two stop codons, two frameshifts), was homologous to VP (36% aa identity with skunk amdoparvovirus VP). This ORF did not possess a conventional Met start codon to express VP1, however, the three aa-long exon leader of a short upstream ORF could potentially provide this, as in other amdoparvoviruses (Qiu et al. 2006). A putative middle (M) protein ORF was identified between the putative NS and VP genes.

The polyglycine (poly-G) region in parvovirus VP proteins is suspected to be responsible for externalizing the VP1u (so that the enzymatic functions of the PLA2 domain can be carried out) as well as exposing the nuclear localization signal (NLS) of the VP1u and the VP2 (M. S. Chapman and Rossmann 1993; Vihinen-Ranta et al. 2002). All exogenous amdoparvovirus VP peptide sequences contain a polyglycine (poly-G), despite lacking a PLA-2 domain. A poly-G region was also present in the predicted VP sequence of ProMuc.1, whereas it was absent from EllLut.1 VP (**Figure S2**). Interestingly, however, the VP1u sequences of both both PVe disclosed a putative NLS. Notably, the ProMuc.1 VP sequence contained numerous indels relative to those of amdoparvoviruses. Notably, however, indels were almost exclusively confined to the VR loops (**Figure 3**). The only insertion, six-aa-long, was present in VR VIII. Interestingly, dependoparvovirus-derived PVe previously identified in marsupial genomes have also been reported to harbor extended VRVII loops (Smith et al. 2016).

**Figure 3.**
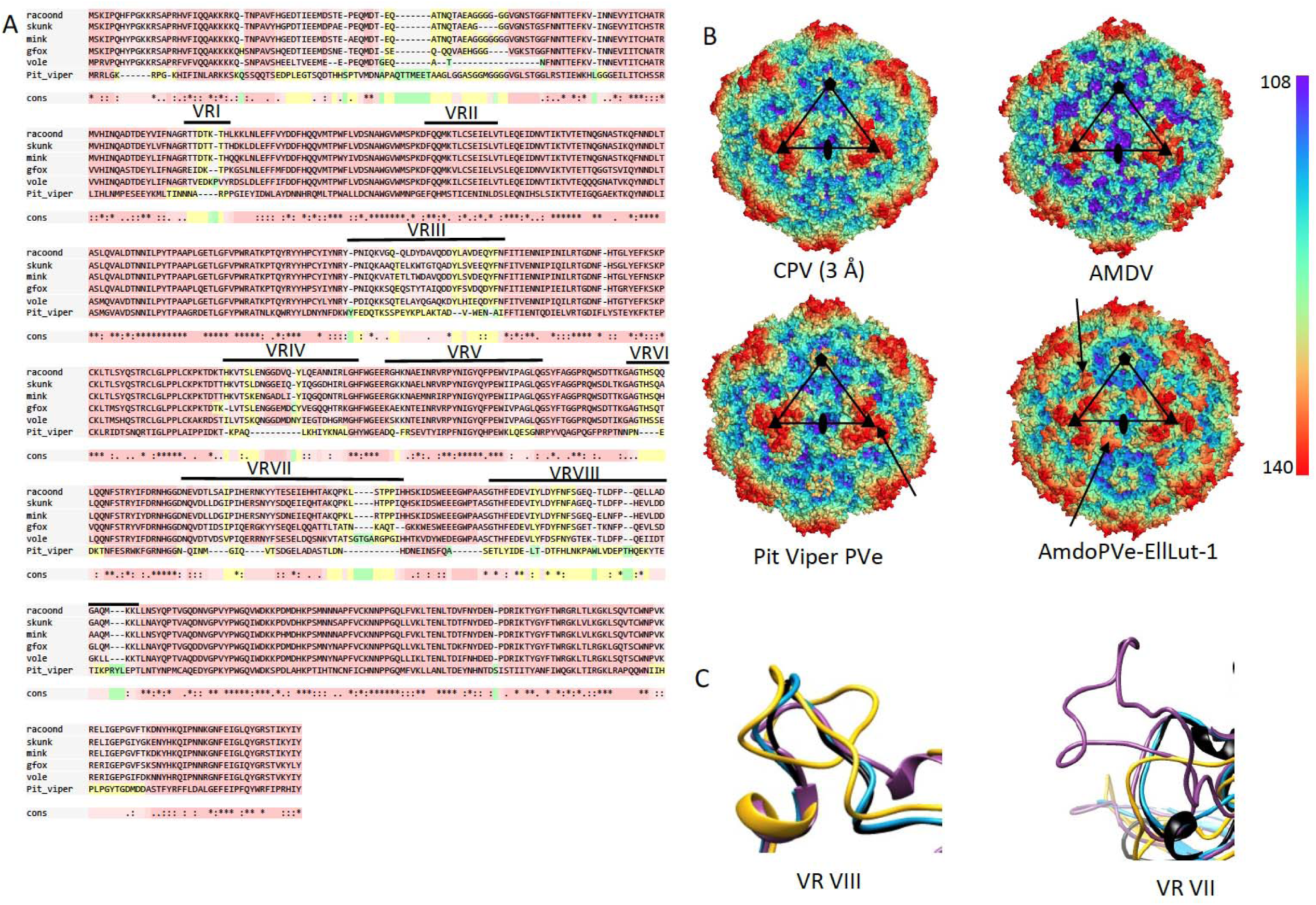
**Panel (a)**: Alignment of the PVe VP protein sequences with those of exogenous amdoparvoviruses. Variable regions (VR) are indicated by horizontal lines. **Panel (b):** Results of homology modeling; the capsid structure of canine parvovirus (CPV) served as a reference structure for all the three further models. The bar shows the distance from the capsid center in Ångströms and the structures are colored accordingly. The pentagon marks the five-fold, the triangles the three-fold and the two-fold is indicated by an ellipse. The arrows mark the VRIII region of ProMuc.1 and the VRVII of the EllLut-1 capsids, which contain the only insertions compared to amdoparvovirus VR regions. (C) Ribbon diagrams of the VRVIII (left) and VRVIII (right) loops of CPV (blue), Aleutian mink disease virus (AMDV) (black), EllLut-1 (pink), and ProMuc.1 (yellow). Abbreviations: racoon=racoondog amdoparvovirus; gfox=grey fox amdoparvovirus; skunk=skunk amdoparvovirus; Mink=Aleutian mink disease virus; vole=Transcaucasian mole vole PVe EilLut.1; Pit viper=pit viper PVe ProMuc.1.

ProMuc.1 occurs on a contig that has not been mapped to a specific chromosome. Nevertheless, the pre-integration locus could be identified in WGS data of two other reptilian species: the Burmese python (*Python bivittatus*), and a colubrid, the common garter snake (*Tamnophis sirtalis*) (data not shown). The absence of a ProMuc.1 insertion in these taxa establishes that it was incorporated into the germline of the pit viper subsequent to its divergence from these species, which is estimated to have occurred 34-54 Mya (Head et al. 2005).

### PVe in mammalian genomes with amdoparvoviral and protoparvoviral characteristics

Genome screening *in silico* identified three additional matches to amdoparvoviruses in mammal genomes (**Table 1**). We examined these Pve and found that all three of were highly fragmented by stop codons, frameshifts and transposable elements. Nevertheless, all three encoded near complete VP peptides, all of which exhibited a well-preserved calcium-binding loop in their N-terminal PLA2 domains (**Figure 1**). However, the catalytic core was barely recognizable in the Cape hyrax element (ProCap.1) and completely absent in the aarvark element (OryAfe.1) (**Figure 1b, Figure S2**). ProCap.1 appeared to lack an NLS sequence, and a poly-G stretch was absent from all three elements, as in the VP encoded by the amdoparvovirus-derived PVe EllLut.1 (**Figure S2**). Apart from the disintegrated catalytic domain of the PLA2, the Tasmanian devil element (SarHar.1) displayed the most well-preserved VP1u sequence.

With the exception of OryAfe.1, which contained a highly disrupted NS homolog spanning 343 aa residues, only a minimal trace of the non-structural genes could be detected (**Figure 1**). In phylogenies based on NS (**Figure 2a**), OryAfe.1 grouped together with the Mpulungu bufavirus of shrews (Sasaki et al. 2015) as a robustly-supported sister group to rodent, ungulate and carnivore protoparvoviruses. In phylogenies based on VP (**Figure 2b**), all three mammal PVe formed a robustly supported clade in a position intermediate between the amdoparvoviruses and protoparvoviruses. The viper element ProMuc.1 grouped basal to this clade, but with weak support.

### Structural characterization of PVe capsid proteins via homology modeling

We investigated the capsid (VP) sequences of the more complete and intact PVe using homology modelling. Using this approach, the capsids encoded by pit viper and mole vole PVe proved to be structurally most similar to the canine parvovirus (CPV) capsid (PDB ID: 2CAS) according to fold recognition, hence this structure was used as a template. As there are currently no publicly available structural data for amdoparvorviruses, we constructed the model of the AMDV capsid as well, based on the CPV template.

The predicted structures of the capsid proteins encoded by the ProMuc.1 and EllLut.1 displayed a rather protoparvovirus-like appearance, unlike the AMDV capsid model (**Figure 3**). In the case of ProMuc.1, three-fold protrusions were thicker and bulkier than either on CPV or AMDV, while the EllLut.1 capsid model displayed spike-like protrusions rather than the slope-like depressions characteristic of the parvovirus two/fivefold wall. These differences could be ascribed to insertions in variable regions, namely VRVIII of the ProMuc.1 and VRVII of EllLut.1. Both capsids appeared to contain the canonical β-strand A (βA), an eight-stranded β-barrel core making up the jelly roll fold ((βBIDG-CHEF), and an α-helix (αA) (**Figure 3b and c**).

## DISCUSSION

### The genomic fossil record of amdoparvoviruses

The assorted PVe sequences being revealed via whole genome sequencing of animal species are a unique and useful source of retrospective information about parvovirus evolution that is in some ways equivalent to a parvovirus ‘fossil record’ (Katzourakis and Gifford 2010). However, while there are eight parvovirus genera currently recognised to infect vertebrates (Cotmore et al. 2014), the PVe that have so far been identified in vertebrate genomes are overwhelmingly derived from two of these genera: *Dependoparvovirus* and *Protoparvovirus.* PVe derived from dependoparvoviruses have been identified in several orders of birds and mammals (Belyi et al. 2010; Cui et al. 2014; Katzourakis and Gifford 2010), while PVe derived from protoparvoviruses have been reported in rodent genomes (Kapoor et al. 2010). Larger numbers of ‘protoparvovirus-like’ PVe are present in the genomes of mammals, including rodents and marsupials (Arriagada and Gifford 2014; Katzourakis and Gifford 2010), but it is less clear whether these derive from *bona fide* protoparvoviruses or a distinct parvovirus lineage (e.g. an extinct genus). Prior to this study, only a single, highly fragmented PVe (ProCap.1) had been reported as showing homology to amdoparvoviruses (Katzourakis and Gifford 2010).

We report the first examples of PVe that are unambiguously derived from amdoparvoviruses. These two elements, which were identified in the genome of the Transcausian mole vole, were found to group within the diversity of amdoparvovirus isolates in molecular phylogenies (**Figure 2**). They also exhibit characteristic features that support their grouping within the genus *Amdoparvovirus*, including the presence of a putative middle (M) ORF, and the absence of the PLA2 domain from the predicted VP protein sequence (**Figure 1, Figure S2**).

We did not identify any other PVe that grouped convincingly within the *Amdoparvovirus* genus. However, we identified several that displayed a mixture of amdoparvoviral and protoparvoviral features. PVe in this ‘amdo-proto’ (AP) group – which may not be monophyletic – grouped in an intermediate position in phylogenetic trees. Furthermore, while these elements appear to be marginally more closely related to protoparvoviruses than to amdoparvoviruses (**Figure 2**), certain aspects of their genome organization suggested an evolutionary connection to amdoparvoviruses. For example, in the pit viper element, these include: (i) the presence of a single promoter and two polyadenylation signals; (ii) the attributes of the intron in the VP1u, and; (iii) the apparent absence of a PLA2 domain (**Figure 1**).

We were able to infer maximum age bounds of 54 million years for the pit viper PVe and 10 million years for the mole vole PVe, based on the identification of empty integration sites in related species. However, none of the PVe reported here were identified as orthologous copies in two or more related species. Consequently, we are unable to draw firm conclusions with respect to their minimum ages. The mutational degradation observed in the more fragmented elements suggests they are likely to have similarly ancient origins to other PVe (i.e. extending back millions of years). In the case of the mole vole, two EllLut.2 alleles were present (one containing a deletion relative to the other), indicating that these elements have likely been present in the species gene pool for multiple generations.

The confirmed host range of amdoparvoviruses is restriced to carnivores, but this likely reflects limited sampling – metagenomic studies of parvovirus diversity are providing strong hints that most if not all genera in the subfamily *Parvovirinae* are likely to have representatives that infect all most if not all extant mammalian orders (de Souza et al. 2018). Nonetheless, the identification of amdoparvovirus-derived PVe in the genome of the Transcaucasian mole vole advances our current knowledge by uneqivocally demonstrating that amdoparvoviruses have infected rodents in the past. The pit viper PVe reported here (ProMuc.1) is the first to be identified in a reptile genome.

### Evolution of amdoparvovirus and protoparvovirus capsid proteins

Comparison of predicted VP protein structures to those of exogenous amdoparvoviruses revealed that most differences are limited to regions of conspicuous functional significance, including the PLA2 domain, VP1u, and the VR loops that are exposed on the virion surface and are thought to be involved in mediating many host-virus interactions, such as immunogenicity and receptor attachment (Huang et al. 2014). The fact that variation is overwhelmingly confined to these regions suggests it largely reflects diversity accumulated through selection on ancestral viruses, rather than mutations acquired post-integration. The strikingly high level of deletions seen in the VR regions of the ProMuc.1 VP protein is consistent with this, since we might expect that the reptilian anti-viral response, in which the adaptive immune system plays relatively small role, would exert selective pressures that were somewhat different to those encountered by parvoviruses infecting mammals (Zimmerman et al. 2010). A similar idea has previously been proposed to explain the characteristically smooth surface features, i.e. shorter or absent VRs, of invertebrate-infecting densoviruses (Simpson et al. 1998).

The highly conserved phospholypase A2 (PLA2) motif in VP is required for escape from the endosomal compartments after entering the host cell (Zadori et al. 2001). Uniquely, amdoparvoviruses lack this domain in their VP1u region (Cotmore et al. 2014; Zadori et al. 2001), and consequently their trafficking is not fully understood. In phylogenies amdoparvoviruses, protoparvoviruses and the AP lineage of PVe generally form a discrete, robustly supported clade, the relationships between which are poorly resolved (**Figure 2**). PVe in the AP lineage are presumably derived from an uncharacterised parvovirus lineage possessing amdoparvovirus and protoparvovirus-like features, but the lack of phylogenetic resolution between the three main clades we observed in phylogenies mean we cannot determine whether PVe in the AP lineage represent transitional forms along a pathway from protoparvovirus-like complete PLA2 domains to the PLA2-absent amdoparvoviral-like VP1u, or an entirely distinct parvovirus lineage. Moreover, the position of the pit viper PVe furtheer complicates this; apart from the possibility of comprising a basal entry of the AP group, it might have evolved independently from these and been member of a yet another separate lineage, making the AP group paraphyletic. We do not imply, however, its direct ancestry to genus *Amdoparvovirus*, but being of a putative fourth lineage, which might be closer related to recent amdoparvoviruses cannot be excluded.

Interestingly, an intact calcium binding loop was found in the predicted VP1u protein sequences of all PVe in the AP lineage that encoded this region, even though most of these sequences also have a degraded PLA2 domain (**Figure 2, Figure S2**). This observation raises the possibility that this loop might have functions in the viral life cycle that are unrelated to its role in phospholipase-mediated escape from the endosomal compartments.

We used homology modelling to infer the structures of the capsid (VP) proteins encoded by PVe. This analysis revealed that both EilLut.1 and ProMuc.1 capsids had a protoparvovirus-like appearance, rather than being similar to the AMDV capsid (**Figure 3**). These findings are intriguing when considered in the light of the phylogenetic relationships depicted in **Figure 2**. Results based on homology modelling should of course be interpreted cautiously, but these observations could reflect that the ancestral viruses that gave rise to these two elements were more similar to protoparvoviruses than amdoparvoviruses in certain aspects of their biology related to their capsid proteins (e.g. tropism or receptor specificity). With regard to this, the prominent role of Fc-receptor-mediated antibody-dependent enhancement (ADE) in AMDV infection should be considered (von Kietzell et al. 2014). Residues 428 to 446 in VP have been shown to play an important role in mediating Fc-receptor attachment during ADE, and interestingly this region overlaps with VR VII, which is highly divergent in both ProMuc.1 and EllLut.1 (**Figure 3**). It is not known whether ADE plays an important role in infections with amdoparvoviruses other than AMDV, or whether the variability observed in this region is relevant to this process, but it is nonetheless intriguing to consider that the distinctive appearance of the AMDV capsid might be related to its use of ADE as an entry mechanism.

### Intact, potentially expressible NS genes encoded by Pve reported here

Two of the PVe described here (EilLut.2 and ProMuc.1) encode intact, expressible NS genes, adding to a growing number of PVe that exhibit this characteristic (Arriagada and Gifford 2014; Katzourakis and Gifford 2010; Liu et al. 2011). Recent studies have shown that independently acquired PVe in rodents and afrotherian genomes exhibit similar patterns of tissue-specific expression in the liver (Arriagada and Gifford 2014; Kobayashi et al. 2018), suggesting that PVe may have been co-opted or exapted by mammalian genomes on more than one occasion. Intriguingly, *in silico* predictions indicated that the intact EilLut.2 replicase identified here could be expressed as a fusion protein with a partial MAFG gene product (**Figure 1a**).

## Acknowledgements

RJG was funded by the Medical Research Council of the United Kingdom (MC_UU_12014/12). JJP and MA-K were supported by a grant from the National Institutes of Health (NIH R01 GM109524). We thank Andrew Davison and Joseph Hughes for their comments and feedback on the manuscript. Data are available in GenBank.

